# Visual-motor integration deficits in 3q29 deletion syndrome

**DOI:** 10.1101/2022.09.15.508134

**Authors:** Rebecca M Pollak, T Lindsey Burrell, Joseph F Cubells, Cheryl Klaiman, Melissa M Murphy, Celine A Saulnier, Elaine F Walker, Stormi Pulver White, Jennifer G Mulle

**Affiliations:** Center for Advanced Biotechnology and Medicine, Robert Wood Johnson Medical School, Rutgers University; Department of Pediatrics, School of Medicine, Emory University; Department of Human Genetics, School of Medicine, Emory University; Department of Psychiatry and Behavioral Science, School of Medicine, Emory University; Marcus Autism Center, Children’s Healthcare of Atlanta and Emory University; Neurodevelopmental Assessment & Consulting Services; Department of Psychology, Emory University; Department of Psychiatry, Robert Wood Johnson Medical School, Rutgers University

**Keywords:** 3q29 deletion, copy number variants, graphomotor weakness, Beery-Buktenica Developmental Test of Visual-Motor Integration, Beery VMI, VMI

## Abstract

**Purpose:** 3q29 deletion syndrome (3q29del) is associated with neuropsychiatric and neurodevelopmental phenotypes. We previously reported that graphomotor weakness is present in up to 78% of individuals with 3q29del. We have now explored nuances of the graphomotor phenotype and its association with other comorbidities in this population.

**Methods:** Participants were recruited from the online 3q29 registry (3q29deletion.org) for two days of deep phenotyping. 32 individuals with 3q29del (62.5% male) were evaluated with the Beery-Buktenica Developmental Test of Visual-Motor Integration (VMI) to assess visual-motor integration. Participants were also evaluated with measures of cognitive ability, executive function, adaptive behavior, and school function.

**Results:** Males with 3q29del performed significantly worse than females on the VMI and Motor Coordination subtest. VMI performance was significantly associated with ADHD diagnosis and cognitive ability. Compared to published data from individuals with 22q11.2 deletion syndrome, individuals with 3q29del showed significantly more impairment.

**Conclusion:** The 3q29 deletion is associated with substantial deficits in visual-motor integration, Visual Perception, and Motor Coordination. Our data suggests that 3q29del may qualify as a nonverbal learning disability, and that all individuals with 3q29del may benefit from early interventions, including occupational therapy.

---

3q29 deletion syndrome (3q29del) is a rare (∼1:30,000) (Kendall et al., 2017; Stefansson et al., 2014) genomic disorder caused by a typically *de novo* 1.6 Mb deletion on the long arm of chromosome 3 (hg19, chr3:195725000–197350000) (Ballif et al., 2008; Glassford et al., 2016; Willatt et al., 2005), encompassing 21 protein-coding genes, three antisense transcripts, one long noncoding RNA, and one microRNA. 3q29del is associated with a wide array of neurodevelopmental and neuropsychiatric phenotypes, including a greater than 40-fold increased risk for schizophrenia (SZ) (Kirov et al., 2012; Marshall et al., 2017; Mulle, 2015; Mulle et al., 2010; Szatkiewicz et al., 2014) and a 19-fold increased risk for autism spectrum disorder (ASD) (Itsara et al., 2009; Pollak et al., 2019; Sanders et al., 2015). Mild to moderate intellectual disability (ID), global developmental delay (GDD), attention deficit/hyperactivity disorder (ADHD), and anxiety are also commonly associated with 3q29del (Ballif et al., 2008; Cox & Butler, 2015; Girirajan et al., 2012; Glassford et al., 2016; Klaiman et al., 2022; Sanchez Russo et al., 2021; Willatt et al., 2005).

Previous work by our team used standardized direct assessment tools to evaluate individuals with 3q29del for cognitive, neurodevelopmental, neuropsychiatric, and somatic phenotypes (Klaiman et al., 2022; Murphy et al., 2018; Sanchez Russo et al., 2021). Individuals with 3q29del had multiple deficits in cognitive and adaptive functioning, high rates of ID and ASD, and clinically significant deficits in executive function (Klaiman et al., 2022; Murphy et al., 2018; Sanchez Russo et al., 2021). Further, an exceptionally high proportion of individuals with 3q29del (78%) qualified for a clinical diagnosis of graphomotor weakness (Sanchez Russo et al., 2021). Graphomotor weakness had not previously been associated with 3q29del, and the nuances of this specific functional deficit remain to be explored.

Graphomotor skills are the foundational fine motor skills required for more advanced activities like writing and drawing (Suggate et al., 2018; Ziviani & Wallen, 2006). Graphomotor skills begin to develop in early childhood with scribbles on paper, and gradually progress to more complex skills like legible handwriting (Ziviani & Wallen, 2006). Graphomotor weakness, in turn, is a clinically significant deficit in these foundational skills. In addition to poor handwriting, graphomotor skills are also strongly associated with the development of basic reading skills (Suggate et al., 2016, 2018) and math skills (Luo et al., 2007). Graphomotor ability can be assessed in early elementary school-aged children (Volman et al., 2006). It has been shown that early intervention can substantially improve graphomotor skills in young children (Ratzon et al., 2007), highlighting the critical importance of identifying graphomotor difficulties early, especially in high-risk populations.

In the present work, we further define the graphomotor phenotype of 3q29del, including possible associations with cognitive, neuropsychiatric, and neurodevelopmental comorbidities within this population. This study provides critical insight into this previously unexplored deficit in 3q29del, with important implications for therapeutic interventions and educational supports, and ultimately contributes to an improved understanding of this patient population.

## Methods

### Study participants

Individuals with 3q29del were recruited from the online 3q29 registry (3q29deletion.org) for 2 days of in-person deep phenotyping, as previously described (Klaiman et al., 2022; Murphy et al., 2018; Sanchez Russo et al., 2021). 32 individuals with 3q29del (62.5% male) were included in the present study, ranging in age from 4.9-39.1 years (mean = 14.5 ± 8.3 years). A description of the study sample can be found in Table 1. This study was approved by Emory University’s Institutional Review Board (IRB00064133) and Rutgers University’s Institutional Review Board (PRO2021001360).

**Table 1.**
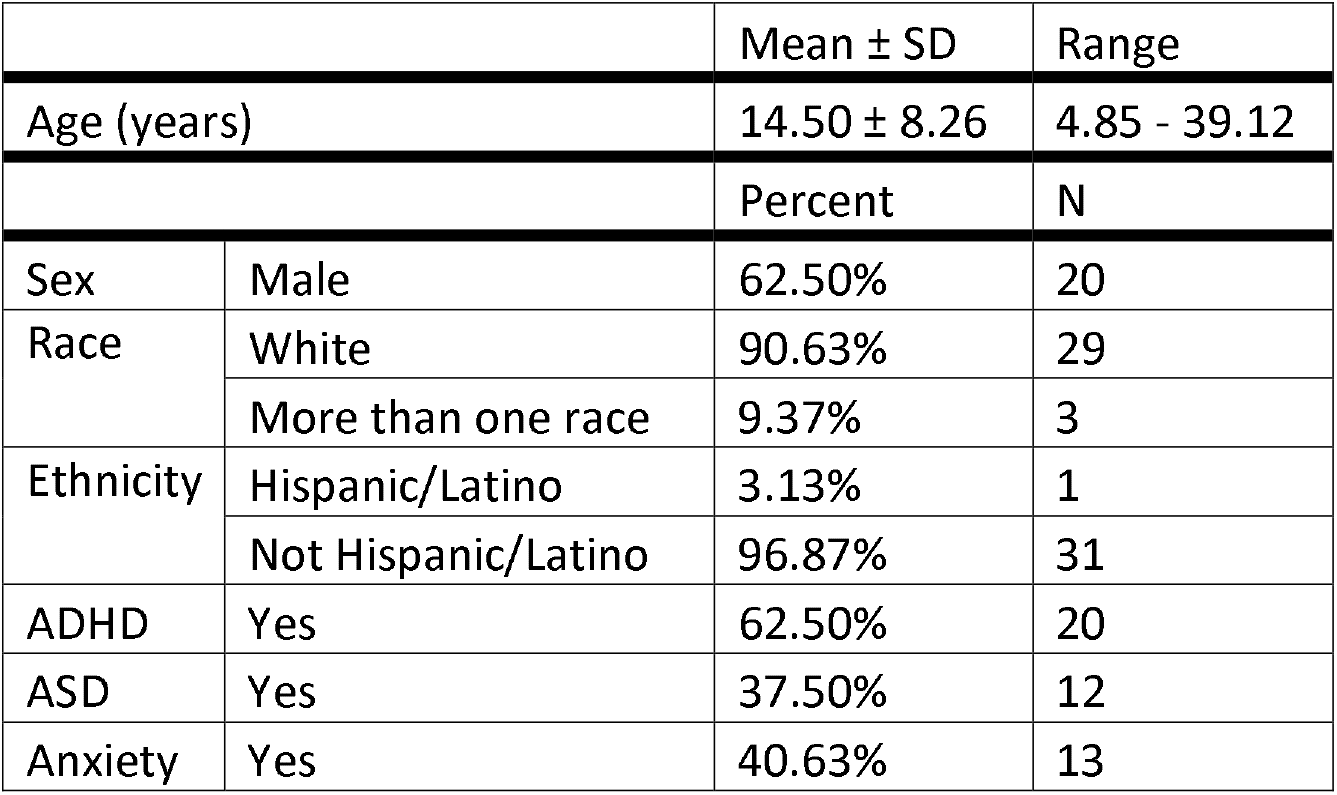
Demographic and clinical data for our 3q29del study sample. Anxiety disorders included generalized anxiety disorder, specific phobia, separation anxiety, and social anxiety disorder.

### Measures

The measures used in this study were as previously described (Klaiman et al., 2022; Murphy et al., 2018; Sanchez Russo et al., 2021). Briefly, visual-motor integration was assessed using the Beery-Buktenica Developmental Test of Visual-Motor Integration, 6^th^ Edition (VMI) (Beery & Beery, 2010), administered by trained PhD-level clinical psychologists. The VMI requires the study participant to copy an increasingly complex series of geometric forms. All study participants were also administered the VMI supplemental subtests to assess Visual Perception and Motor Coordination performance independently (Beery & Beery, 2010). Cognitive ability was evaluated using the Differential Ability Scales, Second Edition (DAS-II) (Elliott et al., 1990) for individuals under 18 years (n = 24) or the Wechsler Abbreviated Scale of Intelligence, Second Edition (WASI-II) (Wechsler, 1999) for individuals 18 years and older (n = 8). Executive function was evaluated using the Behavior Rating Inventory of Executive Function, 2^nd^ Edition (BRIEF-2) for participants 18 years or younger and the Behavior Rating Inventory of Executive Function for Adults (BRIEF-A) for over 18 years (Gioia et al., 2015; Roth & Gioia, 2005). The BRIEF-2 and BRIEF-A (BRIEF) ask the informant to rate the participant’s behaviors associated with nine domains of executive function. The Vineland Adaptive Behavior Scales, Third Edition, Comprehensive Parent/Caregiver Form (Vineland-3) was used to assess adaptive behavior (Sparrow et al., 2016). The Vineland-3 assesses day to day activities in the domains of socialization, daily living skills, communication skills, and motor skills. The school-age Child Behavior Checklist (CBCL) was used to assess school performance for individuals ages 6-18 (n = 19) (Achenbach & Rescorla, 2001). The CBCL assesses behavioral and developmental problems in children between 6 and 18 years of age. Additional detail regarding the assessments used in the present study can be found in the Supplemental Information.

### Analysis

All analyses were performed in R version 4.0.4 (R Core Team, 2008). Standardized scores were used for all analyses. Statistical analysis was performed using simple linear models and generalized linear regression implemented using the stats R package (R Core Team, 2008). Because the standardized VMI scores used in all analyses are age-normed, age was not included as a covariate in any models. Correlation tests were performed using the ppcor R package (Kim, 2015). Comparison of 3q29del to 22q11.2 deletion syndrome was performed using one sample t-tests implemented via the stats R package (R Core Team, 2008). Data visualization was performed using the plotly (Sievert et al., 2017) and ggplot2 (Wickham, 2009) R packages.

## Results

### VMI performance in 3q29del

The VMI is scored on a standard scale, with a mean of 100 and a standard deviation of 15. As previously reported (Sanchez Russo et al., 2021), participants with 3q29del scored on average 2 standard deviations below the population mean of 100 for the VMI (3q29del mean = 69.3 ± 16.7, Figure 1A). Individuals with 3q29del had relatively stronger performance on the Visual Perception subtest, with an average score approximately 1.75 SD below the population mean (3q29del mean = 73.8 ± 17.7, Figure 1A). The most substantial deficit observed in our study population was on the Motor Coordination subtest, with an average score approximately 2.5 SD below the population mean and significantly lower than the respective Visual Perception scores (3q29del mean = 61.3 ± 15.0, p = 0.004; Figure 1A). These data demonstrate that the 3q29 deletion is associated with significant visual-motor integration problems, with the most substantial deficit in Motor Coordination.

**Figure 1.**
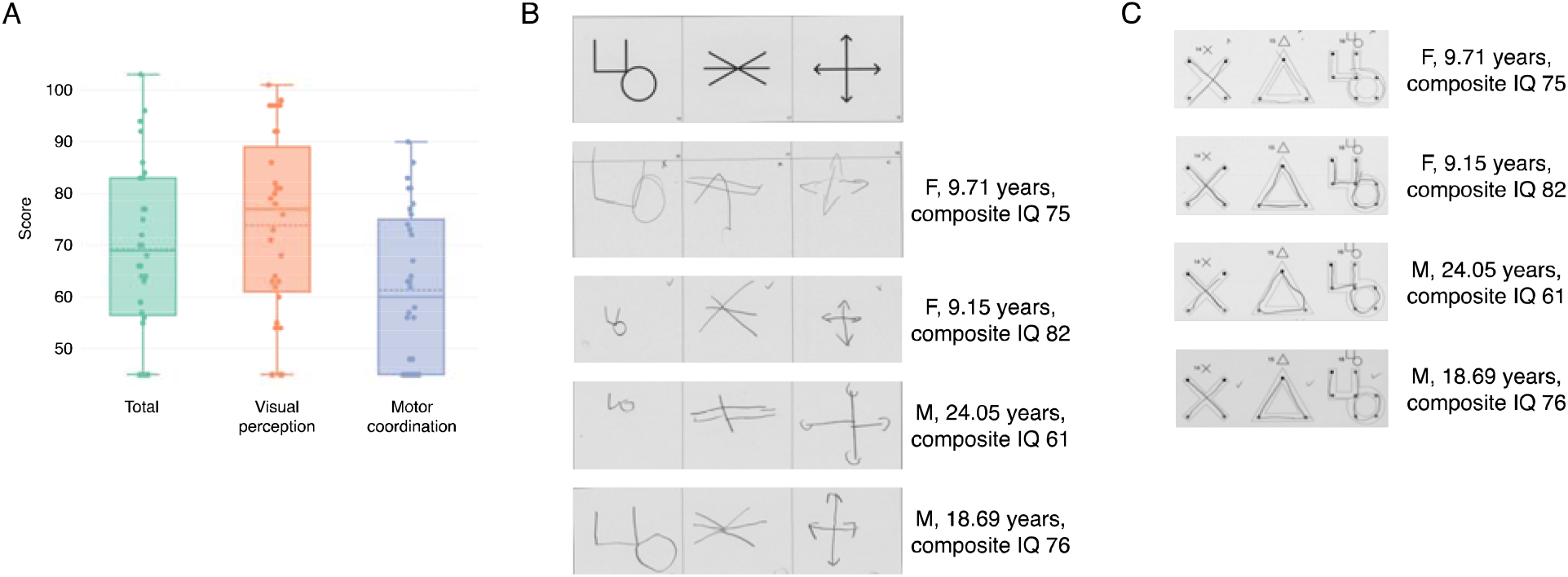
A) Distribution of scores on the VMI, the Visual Perception subtest, and the Motor Coordination subtest for study participants with 3q29del. B) Sample of 3q29del study subject performance on items 16-18 of the VMI. C) Sample of 3q29del study subject performance on items 14-16 of the Motor Coordination subtest.

To better illustrate the range of performance by our 3q29del study subjects, samples of an item from the VMI and an item from the Motor Coordination subtest are shown in Figure 1B-C. The performance varies and is not age-dependent: older participants did not perform any better than younger participants (p > 0.05 for VMI, Visual Perception, and Motor Coordination). Overall, participants with 3q29del struggled on all aspects of the VMI, with only one participant scoring above 100 on the VMI, one participant scoring near 100 on the Visual Perception subtest, and no participants scoring above 100 on the Motor Coordination subtest (Figure 1A).

### Sex-specific differences in VMI performance

There have been sporadic reports of sex differences on the VMI in typically developing children and adults, children with Down syndrome, and children and adults with 22q11.2 deletion syndrome (Coallier et al., 2014; Duijff et al., 2012; Lachance & Mazzocco, 2006; Munro et al., 2012; Niklasson & Gillberg, 2010; Rihtman et al., 2010). To determine whether a sex difference exists in our study population, we performed a sex-stratified analysis of VMI scores. Males with 3q29del scored significantly lower than females with 3q29del on the overall VMI (3q29del male mean = 64.6 ± 15.3, female 3q29del mean = 77.0 ± 16.8, p = 0.04; Figure 2A) and the Motor Coordination subtest (3q29del male mean = 57.6 ± 15.3, 3q29del female mean = 67.6 ± 12.7, p = 0.0005; Figure 2C), indicating significantly greater impairment in males with 3q29del in overall visual-motor integration and in Motor Coordination. There was no statistically significant difference between males and females with 3q29del on the Visual Perception subtest, though males on average scored 12 points below females (3q29del male mean = 69.3 ± 16.8, 3q29del female mean = 81.3 ± 17.4, p > 0.05; Figure 2B). These data show that while both males and females with 3q29del have visual-motor integration deficits, males with 3q29del are more substantially impacted.

**Figure 2.**
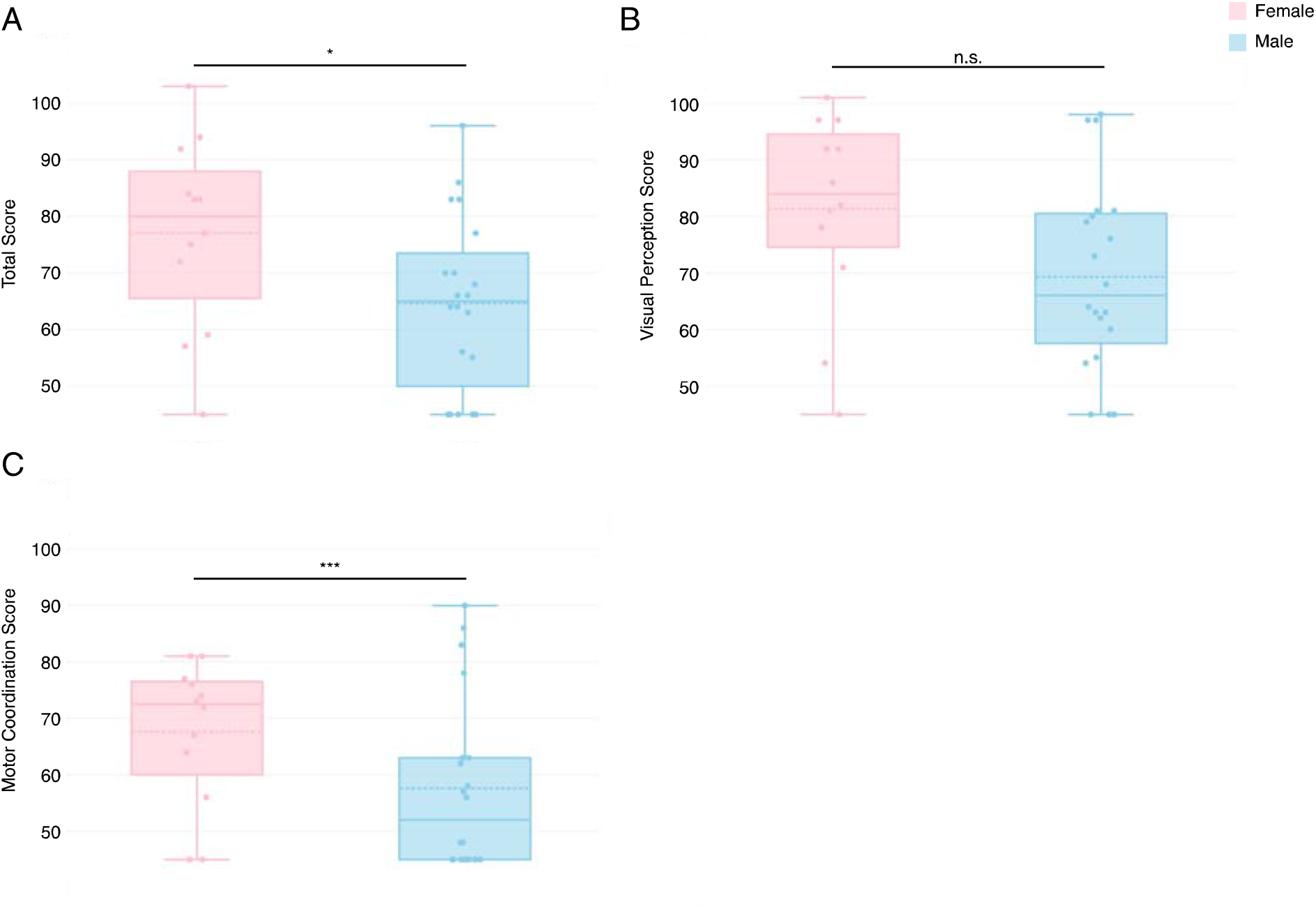
A) VMI score, showing that females with 3q29del (n = 12) score significantly higher than males (n = 20). B) Visual Perception subtest score, showing no significant difference between females and males with 3q29del. C) Motor Coordination subtest score, showing that females with 3q29del score significantly higher than males. n.s., not significant; *, p < 0.05; ***, p < 0.001

### Association between VMI performance and neurodevelopmental and neuropsychiatric phenotypes

There is a well-established association between the 3q29 deletion and a range of neurodevelopmental and neuropsychiatric phenotypes, including ADHD, ASD, and anxiety (Itsara et al., 2009; Kirov et al., 2012; Marshall et al., 2017; Mulle, 2015; Mulle et al., 2010; Pollak et al., 2019; Sanchez Russo et al., 2021; Sanders et al., 2015; Szatkiewicz et al., 2014). To determine whether any of these phenotypes are correlated with overall VMI or subtest performance, we performed stratified analyses. Participants with 3q29del and ADHD (n = 20) scored significantly *higher* on the VMI compared to participants with 3q29del and no ADHD (n = 12) (3q29del+ADHD mean = 73.4 ± 17.2, 3q29del-ADHD mean = 62.4 ± 14.0, p = 0.04; Figure 3A), indicating that clinically significant attentional difficulties are not contributing to the poor performance of 3q29del study subjects on the VMI and suggesting there may be better-preserved visual-motor integration in study subjects with 3q29 del with ADHD. There were no statistically significant differences between participants with 3q29del and ADHD and participants with 3q29del and no ADHD on the Visual Perception (3q29del+ADHD mean = 77.6 ± 16.3, 3q29del-ADHD mean = 67.6 ± 19.0, p > 0.05; Figure S2B) or Motor Coordination subtests (3q29del+ADHD mean = 62.7 ± 16.8, 3q29del-ADHD mean = 59.2 ± 11.7, p > 0.05; Figure S2C), though on average subjects with ADHD scored higher on these subtests. There was no significant association between ASD and VMI score (3q29del+ASD n = 12, 3q29del+ASD mean = 64.3 ± 18.3; 3q29del-ASD n =20, 3q29del-ASD mean = 72.2 ± 15.5; p > 0.05; Figure S2D), Visual Perception score (3q29del+ASD mean = 69.5 ± 21.7, 3q29del-ASD mean = 76.4 ± 14.9, p > 0.05; Figure S2E), or Motor Coordination score (3q29del+ASD mean = 59.8 ± 14.0, 3q29del-ASD mean = 62.3 ± 15.9, p > 0.05; Figure S2F). There was no significant association between anxiety and VMI score (3q29del+anxiety n = 13, 3q29del+anxiety mean = 74.3 ± 15.6; 3q29del-anxiety n = 19, 3q29del-anxiety mean = 65.8 ± 17.0; p > 0.05; Figure S2G), Visual Perception score (3q29del+anxiety mean = 76.9 ± 17.1, 3q29del-anxiety mean = 71.7 ± 18.3, p > 0.05; Figure S2H), or Motor Coordination score (3q29del+anxiety mean = 63.6 ± 15.7, 3q29del-anxiety mean = 59.8 ± 14.7, p > 0.05; Figure S2I). These data suggest that the visual-motor integration deficits associated with 3q29del are not due to these comorbid neurodevelopmental and neuropsychiatric conditions, and are rather an independent phenotype of the deletion.

**Figure 3.**
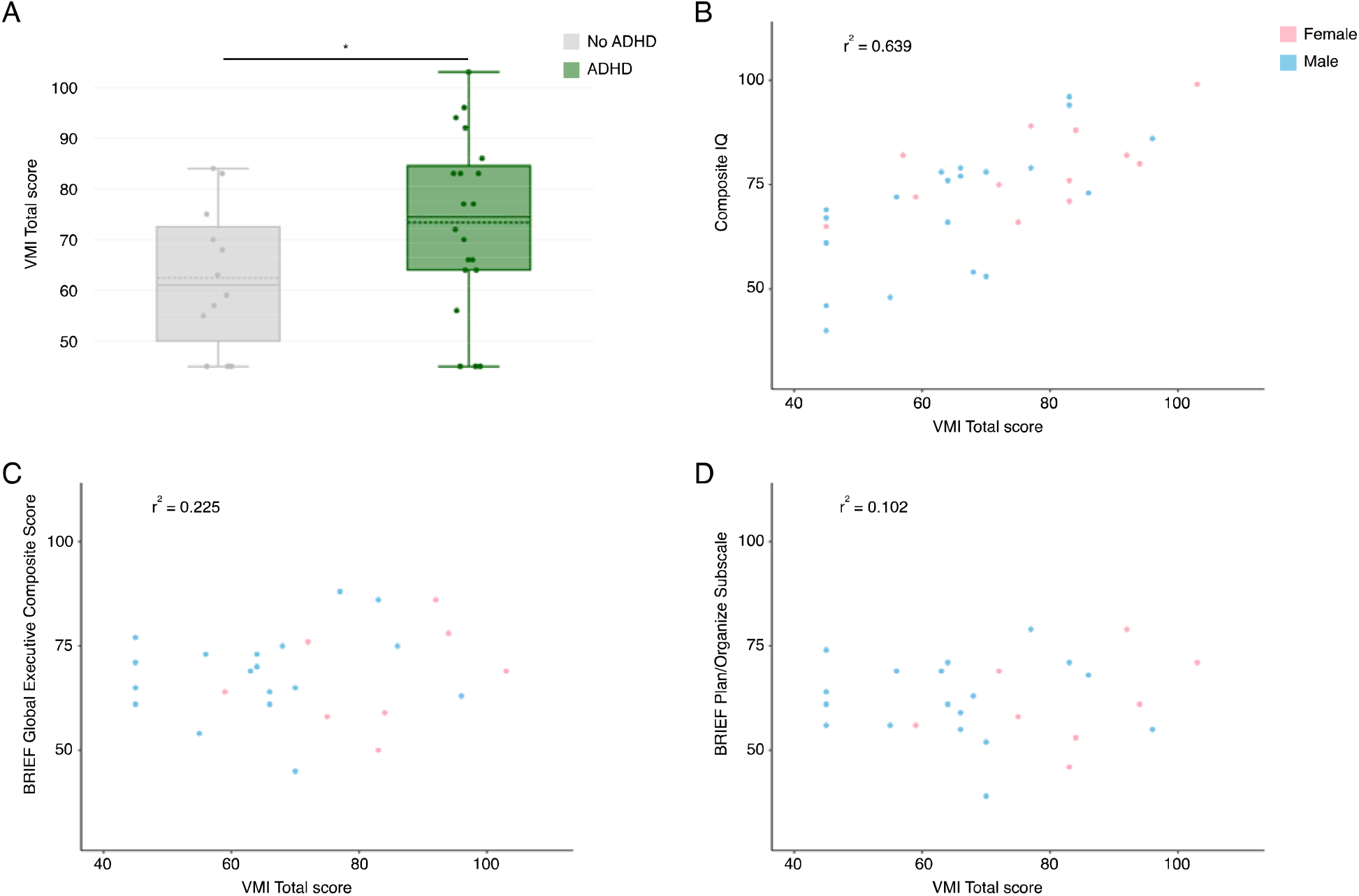
A) VMI score, showing that individuals with 3q29del and ADHD (n = 20) score significantly higher than individuals with 3q29del and no ADHD (n = 12). B) Relationship between VMI score and composite IQ, showing a strong positive correlation. C) Relationship between VMI score and BRIEF Global Executive Composite score, showing a weak positive correlation. D) Relationship between VMI score and BRIEF plan/organize subscale score, showing a very weak positive correlation. *, p < 0.05

### VMI performance is correlated with cognitive ability

Analyses of VMI performance in other genomic disorders have found significant links between VMI score and cognitive ability (Duijff et al., 2012; Heiz & Barisnikov, 2016; Lo et al., 2015; Van Aken et al., 2009). To probe this relationship in our 3q29del study population, we calculated the correlation between VMI performance and measures of cognitive ability. Composite IQ had a strong positive correlation with VMI score (r^2^ = 0.639; Figure 3B), and a moderate positive correlation with the Visual Perception subtest (r^2^ = 0.513; Figure S3B) and the Motor Coordination subtest (r^2^ = 0.550; Figure S3C). Verbal IQ had a weak positive correlation with VMI score (r^2^ = 0.364; Figure S3D), the Visual Perception subtest (r^2^ = 0.226; Figure S3E), and the Motor Coordination subtest (r^2^ = 0.393; Figure S3F). Nonverbal IQ had a strong positive correlation with VMI score (r^2^ = 0.690; Figure S3G) and the Visual Perception subtest (r^2^ = 0.621; Figure S3H), and a moderate positive correlation with the Motor Coordination subtest (r^2^ = 0.428; Figure S3I). Thus, there is a significant relationship between visual-motor integration ability and cognitive ability, but cognitive deficits alone cannot explain the visual-motor integration deficits in 3q29del.

A subset of 24 study subjects under the age of 18 were evaluated with the DAS-II, which includes a measure of spatial ability. Specifically, the Copying and Recall of Designs subtests on the Early Years and School Age batteries, respectively, assess graphomotor and visual-spatial skills similar to the VMI. To determine whether spatial ability was associated with VMI performance in this subset of study subjects, we calculated the correlation between the VMI measures and spatial ability. Spatial ability had a very strong positive correlation with VMI score (r^2^ = 0.805; Figure S4A), a strong positive correlation with the Visual Perception subtest (r^2^ = 0.716; Figure S4B), and a moderate positive correlation with the Motor Coordination subtest (r^2^ = 0.572; Figure S4C). Because overall IQ measures include a component of spatial ability, the correlation between cognitive ability and VMI score may be driven by this measure.

### VMI performance is not related to executive function or daily living skills

A study of VMI performance in children with Williams syndrome motivated the hypothesis that the extremely low VMI scores in Williams syndrome patients were at least partially attributable to executive function challenges, including planning where to place the components of a geometric figure (Bertrand et al., 1997). Williams syndrome has phenotypic similarities to 3q29del, including elevated rates of GDD, ID, and ASD-related symptoms. To determine whether executive function defects are similarly correlated with the poor VMI performance in our 3q29del study subjects, we examined the relationship between VMI performance and the BRIEF. The BRIEF global executive composite score had a weak positive correlation with VMI score (r^2^ = 0.225; Figure 3C), a very weak positive correlation with Visual Perception score (r^2^ = 0.015; Figure S5B), and a weak negative correlation with Motor Coordination score (r^2^ = -0.201; Figure S5C). To determine if the specific planning deficit observed in Williams syndrome is responsible for the poor VMI performance in our 3q29del study subjects, we also examined the plan/organize subscale of the BRIEF. This subscale had a very weak positive correlation with VMI score (r^2^ = 0.102; Figure S5D), almost no correlation with Visual Perception score (r^2^ = -0.008; Figure S5E), and a very weak negative correlation with the Motor Coordination score (r^2^ = -0.112; Figure S5F). This analysis suggests that the poor VMI performance in our 3q29del study subjects is not due to deficits in executive function and may be mechanistically distinct from the VMI deficits in Williams syndrome.

To ask whether graphomotor weakness impacts school performance or activities of daily living, we next tested whether there is a correlation between poor performance on the VMI and deficits in daily living skills or school performance in our 3q29del study subjects. The Vineland-3 Daily Living Skills standard score had a very weak positive correlation with VMI score (r^2^ = 0.174; Figure S6A), and a weak positive correlation with Visual Perception score (r^2^ = 0.230; Figure S6B) and Motor Coordination score (r^2^ = 0.296; Figure S6C). The CBCL school competence score had a weak positive correlation with VMI score (r^2^ = 0.217; Figure S6D), Visual Perception score (r^2^ = 0.299; Figure S6E), and Motor Coordination score (r^2^ = 0.209; Figure S6F). This result implies that intervention to improve visual-motor integration in 3q29del will have a positive but limited impact on daily living skills and school performance.

### Comparing VMI performance in 3q29del and 22q11.2 deletion syndrome

To determine whether the visual-motor integration deficits we observed were unique to 3q29del, we sought to compare results in our study population to previously published VMI performance data from subjects with 22q11.2 deletion syndrome. 22q11.2 deletion syndrome is phenotypically similar to 3q29del, with ID, GDD, and a significantly increased risk for ASD and SZ common between the two syndromes. Studies using the VMI in populations of individuals with 22q11.2 deletion syndrome have reported below average performance on the VMI and the Visual Perception and Motor Coordination subtests (Duijff et al., 2012; Lajiness-O’Neill et al., 2006; Roizen et al., 2010; Van Aken et al., 2009). Using previously published datasets (Duijff et al., 2012; Lajiness-O’Neill et al., 2006; Niklasson & Gillberg, 2010; Roizen et al., 2010; Van Aken et al., 2009), we constructed a 22q11.2 population mean for the VMI and each subtest score and compared these to our 3q29del study subjects. We found that our study subjects with 3q29del scored significantly lower than the constructed 22q11.2 mean on the VMI (3q29del mean = 69.3 ± 16.7, 22q11.2 deletion mean = 87.0 ± 10.4, p = 1.23E-6), the Visual Perception subtest (3q29del mean = 73.8 ± 17.7, 22q11.2 deletion mean = 82.3 ± 15.6, p = 0.01), and the Motor Coordination subtest (3q29del mean = 61.3 ± 15.0, 22q11.2 deletion mean = 76.8 ± 14.3, p = 1.95E-6). This analysis shows that the visual-motor integration deficits in 3q29del are more severe than the deficits identified in 22q11.2 deletion syndrome.

## Discussion

This is the first study to examine aspects of visual-motor integration phenotypes associated with 3q29 deletion syndrome. In previous work by our team, 78% of 3q29del subjects were found to have clinically significant deficits in overall visual-motor integration as assessed by the Beery-Buktenica Developmental Test of Visual-Motor Integration (Sanchez Russo et al., 2021). Here, we explored nuances of that deficit, and found substantial deficits in both visual processing and Motor Coordination. Further, we found no effect of age, but significant sex-dependent differences in performance, with males performing worse than females. VMI performance was significantly associated with ADHD diagnosis, where individuals with 3q29del and ADHD scored *higher* than individuals with 3q29del and no ADHD. Finally, VMI performance was positively correlated with cognitive and spatial ability, but was not associated with executive function, daily living skills, or school performance. VMI performance in 3q29del is considerably worse than what has been reported in 22q11.2 deletion syndrome, despite the similar overall phenotypic profile in both syndromes. VMI performance in 3q29del is also qualitatively different than the deficits reported in Williams syndrome. Taken together, this analysis reveals a unique profile of VMI deficits in 3q29 deletion syndrome.

The 3q29 deletion is associated with a variety of neuropsychiatric and neurodevelopmental phenotypes, including ASD, ADHD, anxiety, and executive function deficits (Itsara et al., 2009; Kirov et al., 2012; Marshall et al., 2017; Mulle, 2015; Mulle et al., 2010; Pollak et al., 2019; Sanchez Russo et al., 2021; Sanders et al., 2015; Szatkiewicz et al., 2014). In the general population, idiopathic ADHD and ASD have independently been associated with significantly worse performance on the VMI (Geurts et al., 2005; Green et al., 2016; Sutton et al., 2011). However, in our study sample ADHD was associated with *better* VMI performance, with individuals with 3q29del and ADHD scoring significantly higher on the VMI. Additionally, ASD and anxiety diagnoses were not associated with VMI performance. We also found that VMI performance was not related to executive function deficits in our study sample. Together, these data indicate that poor visual-motor integration is independent from executive function and attentional deficits within our 3q29del study population. Further studies with larger sample size are needed to better understand the inter- and cross-disorder profiles and if specific syndromic factors play a role in symptom expression.

Mild to moderate ID is observed in the majority of individuals with 3q29del (Ballif et al., 2008; Cox & Butler, 2015; Girirajan et al., 2012; Glassford et al., 2016; Klaiman et al., 2022; Sanchez Russo et al., 2021; Willatt et al., 2005), and previous studies have shown that low IQ can contribute to poor VMI performance in other genomic disorders (Bertrand et al., 1997; Duijff et al., 2012; Lo et al., 2015; Van Aken et al., 2009; Wang et al., 1995). Indeed, composite IQ, verbal IQ, nonverbal IQ, and spatial ability were all positively associated with VMI performance in our study sample. However, none of these variables were perfectly correlated with VMI performance, which suggests that ID alone cannot fully explain the visual-motor integration or motor coordination deficits observed in our study population. Studies of VMI performance in other genomic disorders have found that children with 22q11.2 deletion syndrome or Williams syndrome perform significantly worse than IQ-matched controls (Bertrand et al., 1997; Van Aken et al., 2009). Another study compared VMI performance between individuals with Williams syndrome and Down syndrome and found that the individuals with Williams syndrome had a significantly lower mean score (Wang et al., 1995). These studies indicate that the genomic insults in Williams syndrome and 22q11.2 deletion syndrome impact visual-motor integration ability above and beyond the insult to cognitive ability; the present study suggests the same may be true in the context of the 3q29 deletion. Indeed, previous work by our team has identified a significant association between VMI performance and cerebellar white matter volume (Sefik et al., 2022), suggesting that VMI performance in 3q29del is related to a brain region distinct from those canonically involved in cognition, but likely to contribute to visuo-spatial function.

Poor VMI performance has been associated with deficits in fine motor skills and school problems due to difficulties with handwriting and other critical classroom skills (Daly et al., 2003; KIlIçöz et al., 2022; Sortor & Kulp, 2003; Weil & Amundson, 1994). We found that individuals with 3q29del scored lowest on the Motor Coordination subtest, with multiple individuals earning the lowest possible standardized score. Case studies of 3q29del have reported some gross motor phenotypes, including gait abnormalities (Cox & Butler, 2015), and previous work by our team identified mild to moderate fine motor challenges associated with the 3q29 deletion (Sanchez Russo et al., 2021). However, these challenges do not appear to drive poor VMI performance in our study population (Table S1). Additionally, VMI performance was not related to daily living skills measured by the Vineland-3 or school performance measured by the CBCL. It is possible that visual-motor integration deficits in 3q29del are truly not associated with challenges in daily living skills or school performance, but it is also possible that the measures we used did not adequately capture the challenges that individuals with 3q29del experience in these arenas.

We identified a significant sex difference in our 3q29del study population, with males performing worse than females on the VMI, the Visual Perception subtest, and the Motor Coordination subtest. Additionally, consistent with prior findings in the general population (Maccoby & Jacklin, 1972), females with 3q29del have a higher mean verbal IQ compared to males (female 3q29del mean = 86.33 ± 10.25, male 3q29del mean = 75.55 ± 23.19, p = 0.0009; Figure S7). However, the weak correlation between verbal IQ and VMI performance suggests that this difference is not driving sex differences in VMI performance; rather, sex differences in VMI performance in our 3q29del study subjects appear to be an independent phenomenon. Sex differences in VMI performance have been sporadically reported in the literature, both in studies of individuals without a genomic disorder and in studies of individuals with Down syndrome and 22q11.2 deletion syndrome (Coallier et al., 2014; Duijff et al., 2012; Lachance & Mazzocco, 2006; Munro et al., 2012; Niklasson & Gillberg, 2010; Rihtman et al., 2010). Studies of children in the general population have identified better performance in females relative to males (Coallier et al., 2014; Lachance & Mazzocco, 2006), and studies of adults in the general population have found the reverse, with males performing significantly better than females (Munro et al., 2012). Studies of Down syndrome and 22q11.2 deletion syndrome that have examined sex differences in VMI performance have found that females perform significantly better than males (Duijff et al., 2012; Niklasson & Gillberg, 2010; Rihtman et al., 2010); however, the samples were overwhelmingly children. It is unclear whether the change in relative performance with increasing age, observed in some general population samples, would also occur in populations with genomic disorders. The sample size in the current study is not large enough to interrogate sex by age interactions; future studies with larger sample sizes will be needed to explore these relationships in the 3q29del population.

There are several aspects of VMI performance that are similar between our 3q29del study sample and reports on other genomic disorders. Visual-motor integration deficits have been identified in individuals with 22q11.2 deletion syndrome (Duijff et al., 2012; Lajiness-O’Neill et al., 2006; Niklasson & Gillberg, 2010; Roizen et al., 2010; Van Aken et al., 2009), Prader-Willi syndrome (Dykens, 2002; Lo et al., 2015), and Williams syndrome (Bellugi, 1988; Bertrand et al., 1997; Dykens et al., 2001; Fu et al., 2015; Heiz & Barisnikov, 2016; Pagon et al., 1987; Pezzini et al., 1999; Wang et al., 1995). While the degree of deficit varies between genomic syndromes, there are some qualitative similarities with 3q29del. Similar to our findings in 3q29del, a significant sex difference has been reported in a largely pediatric sample of individuals with 22q11.2 deletion syndrome, with females outperforming males (Duijff et al., 2012; Niklasson & Gillberg, 2010). In Williams syndrome, published images of VMI response items (Bertrand et al., 1997; Wang et al., 1995) bear striking similarity to what we observe in 3q29del. However, we were unable to statistically compare the performance of individuals with Williams syndrome to our data on 3q29del because these studies report raw scores, rather than age-normed standardized scores.

In addition to the qualitative similarities in VMI performance between 3q29del and 22q11.2 deletion syndrome and Williams syndrome, there are also cognitive similarities between these syndromes. 22q11.2 deletion syndrome and Williams syndrome have both been associated with nonverbal learning disabilities (NLDs) (Don et al., 1999; Goldberg et al., 1993; Lepach & Petermann, 2011; MacDonald & Roy, 1988; Niklasson et al., 2001; Rourke, 1995; Rourke et al., 2002; Shprintzen et al., 1978; Swillen et al., 1997; Swillen et al., 1999). NLDs are characterized by deficits in visual-spatial-organizational ability, perceptual deficits, psychomotor impairments, and a difference between verbal and nonverbal IQ (Harnadek & Rourke, 1994; Mammarella & Cornoldi, 2014; Myklebust, 1975; Rourke, 1988; Rourke et al., 2002; Tsur et al., 1995). Previous work by our team identified a significant performance gap between verbal and nonverbal IQ, with verbal IQ substantially higher than nonverbal (Klaiman et al., 2022). Taken together with the substantial visual-motor integration deficits we observed in our 3q29del study sample, these data suggest that 3q29del may also qualify as an NLD. NLDs are a unique class of learning disabilities because they are not typically associated with extremely poor school performance (Mammarella & Cornoldi, 2014), and can be difficult to identify given the well-preserved verbal ability in affected individuals. Approaching 3q29 deletion syndrome as an NLD may further help to shape management strategies that directly address the most fundamental deficits of the syndrome in a cohesive way, thus maximizing long-term benefits for this community.

There are notable qualitative similarities in graphomotor function between 3q29del and other genomic syndromes, but these deficits may arise from distinct mechanisms. A study of individuals with Williams syndrome hypothesized that poor VMI performance is due to a failure to plan how to execute a given drawing, particularly with respect to the placement of components from complex geometric forms in relation to one another (Bertrand et al., 1997). However, we did not find any meaningful association between VMI performance and executive function in our 3q29del study subjects. We also specifically examined the plan/organize from the BRIEF and did not find an association with VMI performance. Mechanisms contributing to poor visual-motor integration in other genomic disorders are not well-described, but it is possible that syndrome-specific cognitive insults are driving the ultimate graphomotor weakness phenotype across syndromes.

While this is the first study to assess detailed visual-motor integration phenotypes in 3q29del, it is not without limitations. First, due to our small sample size we were underpowered for some group comparisons. Additionally, we were unable to assess the effect of race and ethnicity because our sample is overwhelmingly white and non-Hispanic. Efforts are underway to expand recruitment in historically underserved communities, which will enable us to increase our sample size and diversity in future studies. Second, the standardized score range for the VMI has a lower limit of 45. Several of our study subjects scored a 45 on the VMI, the Visual Perception subtest, and/or the Motor Coordination subtest. Thus, we may not be capturing the true lower extent of the range for all three assessments used in the present study. Studies using the VMI in Williams syndrome typically use raw scores rather than standardized scores for this reason; future analyses of this population may be served by examining raw scores in addition to standardized scores, especially for individuals falling in the lower tail of the distribution.

In summary, this is the first study to examine nuances of visual-motor integration ability in individuals with 3q29del. We find substantial deficits in overall visual-motor integration, as well as in visual perception and motor coordination skills, with the greatest deficits in motor coordination. Notably, while these deficits have some correlation with IQ, they appear to be a unique set of phenotypes rather than simply a derivative measure of cognitive disability. Additionally, poor performance on the VMI is independent from ASD, anxiety, and executive function deficits in this sample. The data presented here, in combination with our previous work, suggest that 3q29del may qualify as an NLD. Deficits in nonverbal skills, including nonverbal IQ and visual-motor integration, have not previously been fully appreciated as fundamental deficits associated with 3q29 deletion syndrome, and may have substantial functional impacts on affected individuals and their families. The results of this study suggest that tailored interventions including occupational therapy to address graphomotor skills may improve outcomes for individuals with 3q29 deletion syndrome. Additionally, the pervasive nature of visual-motor integration deficits may offer clues about the biological mechanism underlying the pathology in 3q29 deletion syndrome.

## Declarations

### Ethics approval and consent to participate

This study was approved by Emory University’s Institutional Review Board (IRB00064133) and Rutgers University’s Institutional Review Board (PRO2021001360).

### Consent to participate

All study subjects gave informed consent prior to participating in this study.

### Consent to publish

Not applicable.

### Welfare of animals

Not applicable.

### Availability of data and materials

The datasets used and/or analyzed during the current study are available from the corresponding author on reasonable request and via NDAR (Collection #2614).

### Author’s contributions

RMP performed the statistical analysis, produced all figures and tables, and wrote the manuscript. TLB, JFC, CK, CAS, EFW, and SPW performed the clinical evaluations of 3q29 deletion study participants. TLB, JFC, CK, MMM, CAS, EFW, and SPW helped with data interpretation. JGM edited the manuscript and provided guidance on analyzing and interpreting data. JGM was the principal investigator responsible for study direction. All authors participated in commenting on the drafts and have read and approved the final manuscript.

## Acknowledgements

We gratefully acknowledge our study population, the 3q29 deletion community, for their participation and commitment to research. funding agencies and grant numbers. This study was funded by the National Institute of Mental Health grant R01 MH110701 (PI Mulle).

