## Supplemental Information for "Visual-motor integration deficits in 3q29 deletion syndrome"

**Article title:** Visual-motor integration deficits in 3q29 deletion syndrome

**Journal name:** Journal of Autism and Developmental Disorders

**Supplemental Methods**

*VMI*

The VMI requires the participant to copy a series of geometric forms arranged in order of increasing difficulty, from lines and simple shapes to complex geometric forms. Each page of the VMI test booklet has six blocks, arranged in two rows of three blocks each. The participant is instructed to examine the shape in the top block and reproduce it in the empty block below. To complete the task, the participant is given a pencil with no eraser. The VMI is not a timed assessment. Each item on the VMI is scored 1 or 0, according to whether the participant correctly replicated the shape. Once a participant scores 0 for three consecutive items, scoring is complete. All participants also completed the supplemental VMI subtests, which assess motor coordination and visual perception independently. The Motor Coordination subtest requires the participant to trace a series of increasingly complex geometric forms while staying within guidelines. The Visual Perception subtest requires the participant to correctly identify a matching form in a list and has no graphomotor demands. There is a 5 minute time limit for both the Motor Coordination and Visual Perception subtests.

For analysis, age-normed standardized scores were used. Score conversion was performed according to the supplemental norms (1). Standardized scores have a mean of 100 and a standard deviation of 15. Higher scores on the VMI indicate stronger visual-motor integration.

*DAS-II*

The DAS-II is a measure of cognitive ability designed for children ages 2 years 6 months to 17 years 11 months. There are three separate subtests that assess verbal, nonverbal, and spatial ability. A standardized General Conceptual Ability composite score is also generated to reflect overall cognitive ability. There are two versions of the DAS-II that are administered according to the age of the participant: the DAS-II Early Years Battery is administered to individuals 6 years of age and younger, and the DAS-II School Age Battery is administered to individuals ages 7 to 18. The DAS-II Early Years Battery has two versions, the Lower Early Years and the Upper Early Years, that were administered based on age and developmental ability. In the present study, seven participants were administered the DAS-II Early Years Battery (six participants 6 years of age or younger and one 11 year old participant with extremely limited verbal ability), and 17 participants were administered the DAS-II School Age Battery.

For analysis, age-normed standardized scores were used. Score conversion was performed according to the companion algorithm (2). Standardized scores have a mean of 100 and a standard deviation of 15. Higher scores on the DAS-II indicate less impaired cognitive ability.

*WASI-II*

The WASI-II is a measure of cognitive function for individuals age 6 to 90 years. The WASI-II is a brief measure of cognitive ability, but the scores generated by the assessment are reliable (3). The WASI-II estimates verbal and nonverbal cognitive ability, and a full scale IQ is generated as a measure of overall cognitive ability. In the present study, eight participants 18 years of age or older were administered the WASI-II.

For analysis, age-normed standardized scores were used. Score conversion was performed according to the companion algorithm (3). Standardized scores have a mean of 100 and a standard deviation of 15. Higher scores on the WASI-II indicate less impaired cognitive ability.

*BRIEF*

The BRIEF is a measure of executive function. For participants up to 18 years of age, the parent/informant forms (BRIEF-2) were completed by the parent or guardian, and for participants over 18 years of age the Adult Version forms (BRIEF-A) were completed by the participant. During the assessment, the informant is asked to rate the participant’s behaviors associated with nine dimensions of executive function: inhibiting distractions, self-monitoring, shifting, emotional control, initiation, working memory, planning, organization, and task monitoring. In the present study, the BRIEF was completed using the publisher’s website.

For analysis, age-normed scores were used. Scores were calculated using the publisher’s website (PARiConnect). Higher scores on the BREIF indicate more impaired executive function.

*Vineland-3*

The Vineland-3 is a measure of overall adaptive behavior. The Vineland-3 assesses day to day activities in the domains of socialization, daily living skills, communication skills, and motor skills. Scores from each domain are used to calculate an Adaptive Behavior Composite score, which is an overall estimate of the participant’s adaptive function. In the present study, the Comprehensive Parent/Caregiver Form was completed by a parent or guardian using the publisher’s website.

For analysis, age-normed scores were used. Scores were calculated using the publisher’s web tool (Pearson q-Global). Higher scores on the Vineland-3 indicate less impaired adaptive skills.

*CBCL*

The school-age CBCL is a 113-item, 3-point Likert-scaled questionnaire designed to assess behavioral and developmental problems in children between 6 and 18 years of age (4). The CBCL is completed by the parent or guardian of the participant. The items on the CBCL are grouped into empirically based syndrome scales designed via factor analysis, and DSM-oriented scales designed by expert clinicians. The CBCL also includes competence scale scores, which assess activities, social, and school function.

For analysis, age- and sex-normed scores were used. Score conversion was performed according to the companion algorithm (4). The CBCL school competence score was used for analysis. Higher scores on the CBCL school competence scale less impaired school performance.

**Table S1. Correlation between VMI performance and clinical variables.**

| Scale | Total VMI correlation | Visual perception correlation | Motor coordination correlation |
| --- | --- | --- | --- |
| Vineland-3 Adaptive Behavior | 0.158 | 0.183 | 0.185 |
| Vineland-3 Gross Motor | -0.279 | 0.024 | -0.071 |
| Vineland-3 Fine Motor | -0.071 | 0.213 | 0.178 |
| Vineland-3 Motor Domain | -0.219 | 0.091 | -0.033 |
| BRIEF Shift | 0.033 | -0.140 | -0.278 |
| BRIEF Task Monitor | -0.028 | -0.106 | -0.118 |
| BRIEF Working Memory | 0.173 | -0.007 | -0.080 |
| BRIEF Initiate | 0.118 | 0.024 | -0.134 |
| BRIEF Emotional Control | 0.265 | 0.040 | -0.218 |
| BRIEF Self Monitor | 0.151 | 0.063 | -0.255 |
| BRIEF Inhibit | 0.164 | -0.060 | -0.203 |
| BRIEF Organization of Materials | 0.211 | 0.126 | -0.066 |

Correlation between VMI performance and Vineland-3 and BRIEF variables.

**Figure S1. VMI performance by study subjects with 3q29del.** A) Sample of performance on the VMI by study subjects with 3q29del. B) Performance on an earlier item of the VMI by two 3q29del study subjects that were unable to complete the simple shapes section of the assessment. C) Sample of performance on the motor coordination subtest by study subjects with 3q29del.

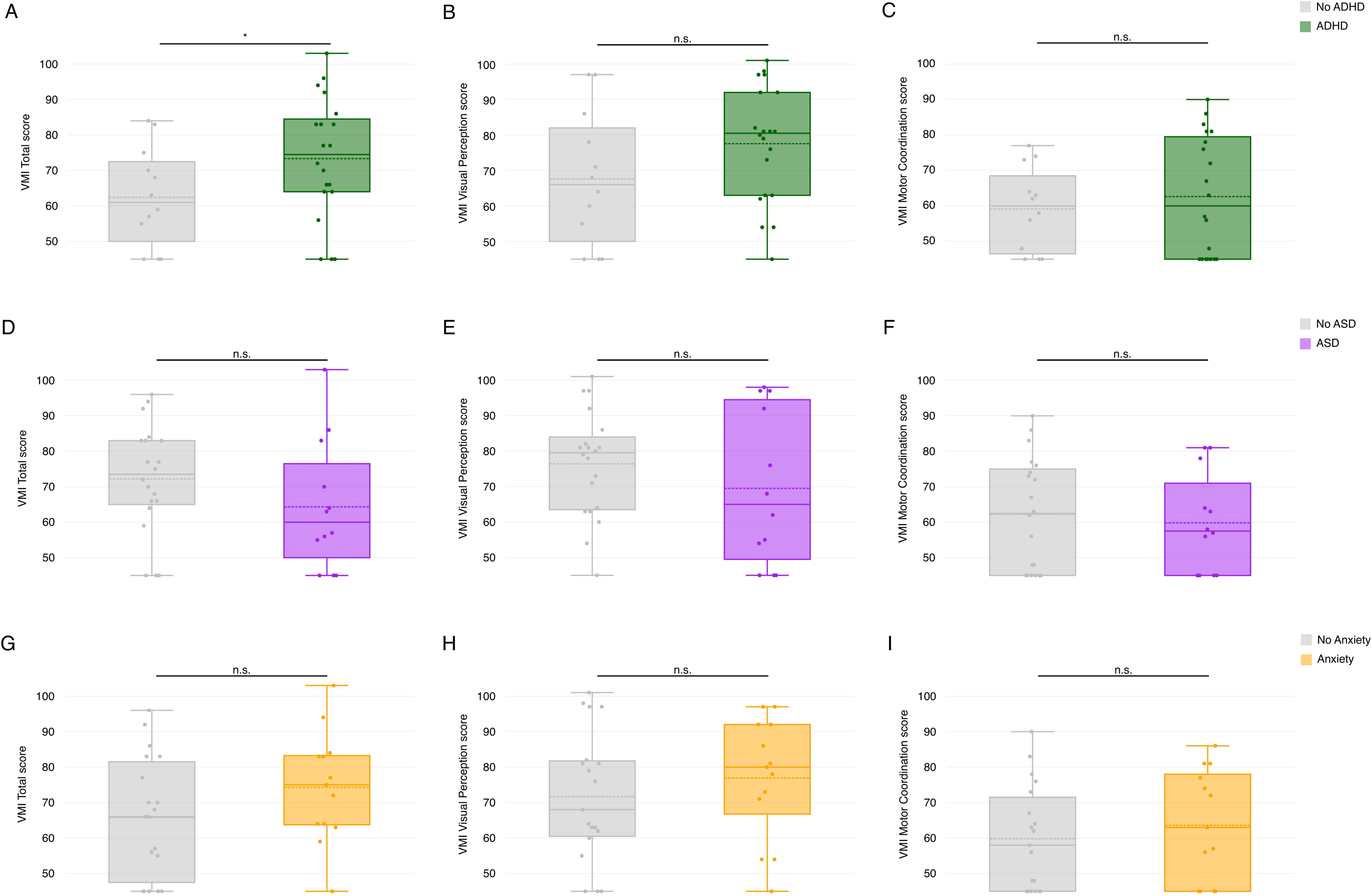

**Figure S2. Association of VMI performance with clinical diagnoses.** A) VMI score, showing that individuals with 3q29del and ADHD (n = 20) score significantly higher than individuals with 3q29del and no ADHD (n = 12). B) Visual perception subtest score and C) motor coordination subtest score, showing no significant difference between individuals with 3q29del and ADHD and individuals with 3q29del and no ADHD. D) VMI score, E) visual perception subtest score, and F) motor coordination subtest score, showing no significant difference between individuals with 3q29del and ASD (n = 12) and individuals with 3q29del and no ASD (n = 20). G) VMI score, H) visual perception subtest score, and I) motor coordination subtest score, showing no significant difference between individuals with 3q29del and anxiety (n = 13) and individuals with 3q29del and no anxiety (n = 19).

n.s., not significant; *, p < 0.05

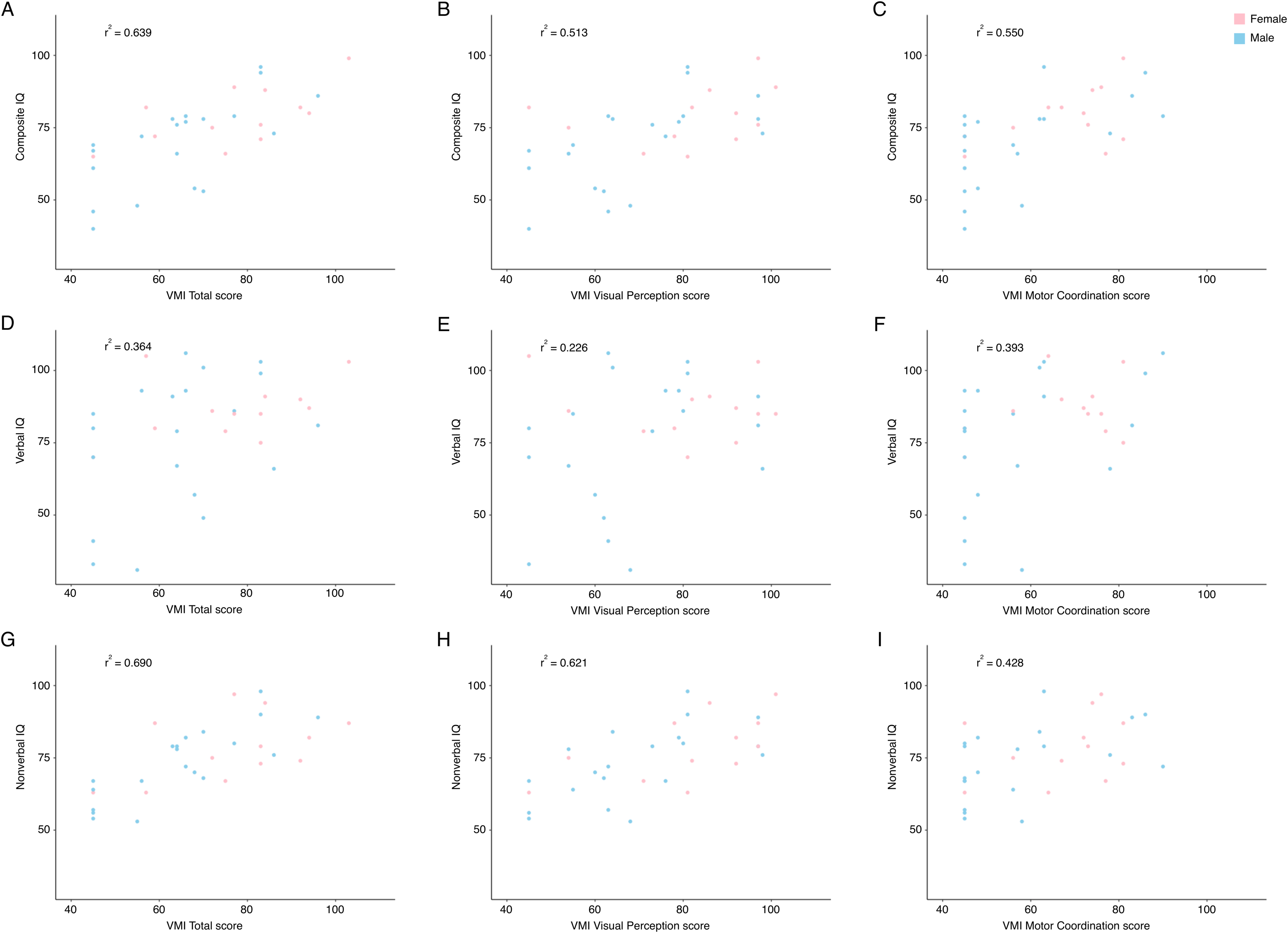

**Figure S3. Association of VMI performance and IQ scores.** A) Relationship between VMI score and composite IQ, showing a strong positive relationship. B) Relationship between the visual perception subtest score and composite IQ, showing a moderate positive relationship. C) Relationship between the motor coordination subtest score and composite IQ, showing a moderate positive relationship. D) Relationship between VMI score and verbal IQ, showing a weak positive relationship. E) Relationship between the visual perception subtest score and verbal IQ, showing a weak positive relationship. F) Relationship between the motor coordination subtest score and verbal IQ, showing a weak positive relationship. G) Relationship between VMI score and nonverbal IQ, showing a strong positive relationship. H) Relationship between the visual perception subtest score and nonverbal IQ, showing a strong positive relationship. I) Relationship between the motor coordination subtest score and nonverbal IQ, showing a moderate positive relationship.

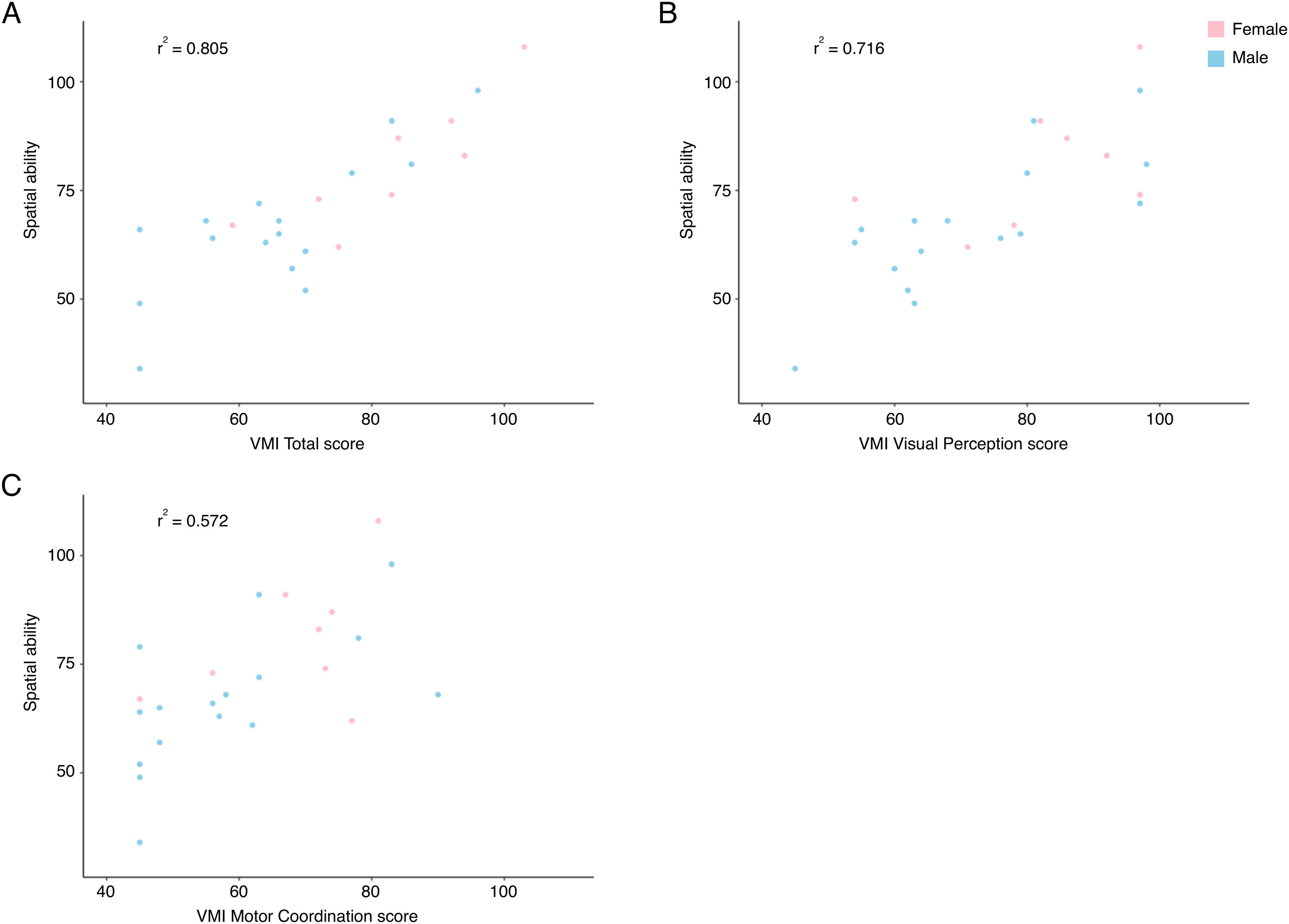

**Figure S4. Association of VMI performance and spatial ability.** A) Relationship between VMI score and DAS-II spatial ability, showing a very strong positive relationship. B) Relationship between the visual perception subtest score and DAS-II spatial ability, showing a strong positive relationship. C) Relationship between the motor coordination subtest score and DAS-II spatial ability, showing a moderate positive relationship.

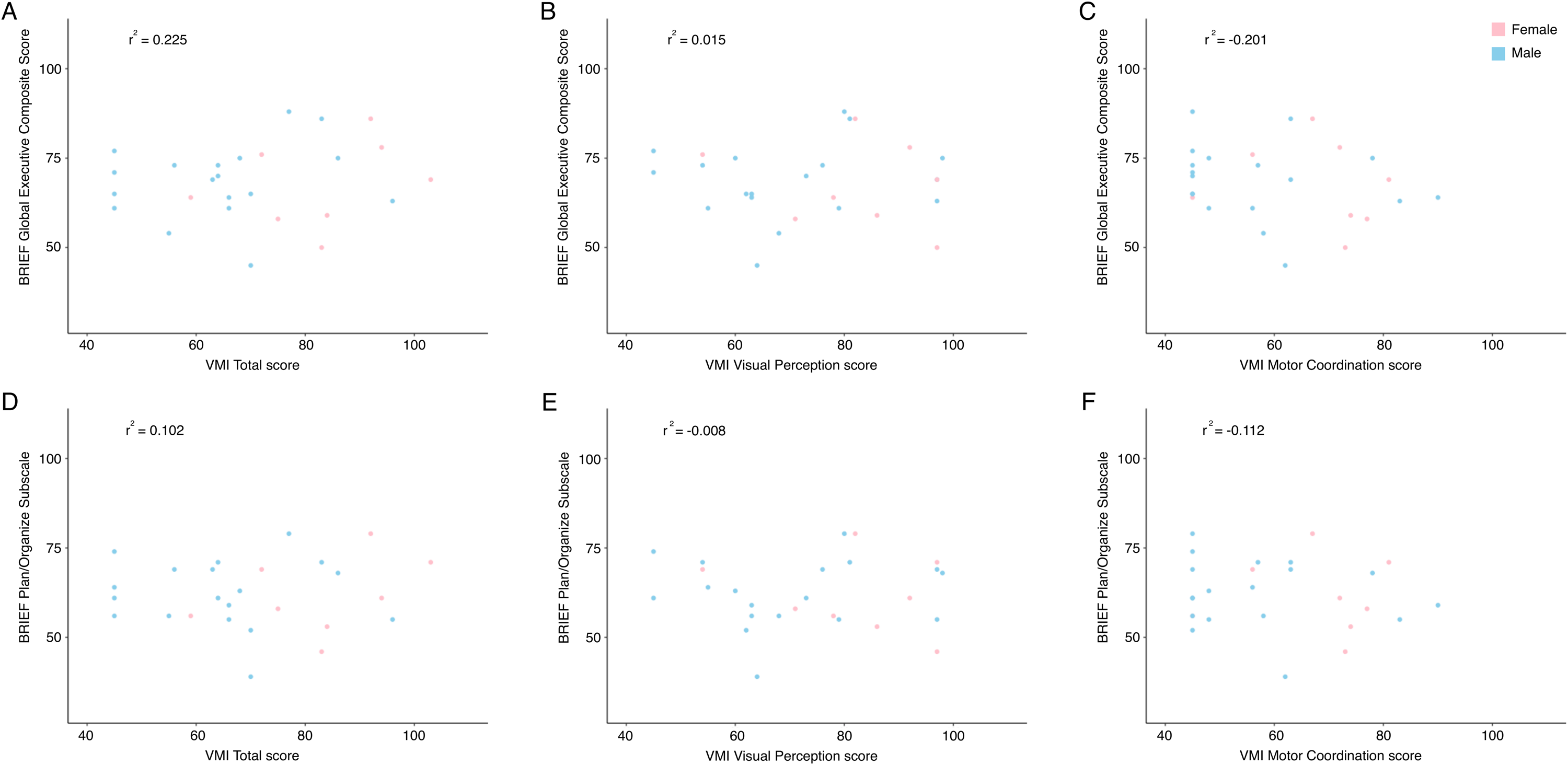

**Figure S5. Association of VMI performance and executive function.** A) Relationship between VMI score and the BRIEF Global Executive Composite score, showing a weak positive relationship. B) Relationship between the visual perception subtest score and the BRIEF Global Executive Composite score, showing a very weak positive relationship. C) Relationship between the motor coordination subtest score and the BRIEF Global Executive Composite score, showing a weak negative relationship. D) Relationship between VMI score and the BRIEF plan/organize subscale score, showing a very weak relationship. E) Relationship between the visual perception subtest score and the BRIEF plan/organize subscale score, showing a very weak negative relationship. F) Relationship between the motor coordination subtest score and the BRIEF plan/organize subscale score, showing a very weak negative relationship.

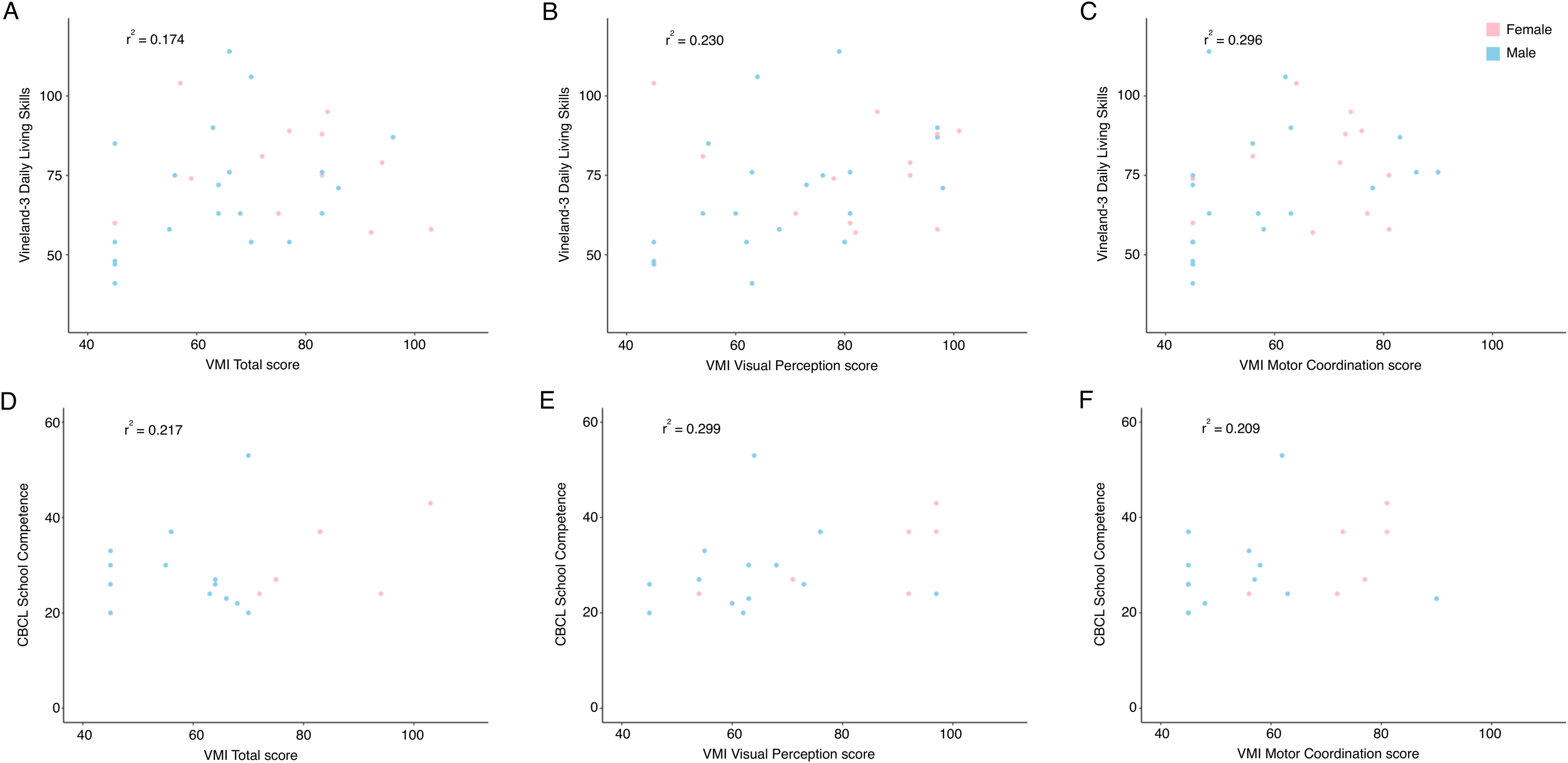

**Figure S6. Association of VMI performance and daily living skills and school performance.** A) Relationship between VMI score and the Vineland-3 Daily Living Skills standard score, showing a very weak positive relationship. B) Relationship between the visual perception subtest score and the Vineland-3 Daily Living Skills standard score, showing a weak positive relationship. C) Relationship between the motor coordination subtest score and the Vineland-3 Daily Living Skills standard score, showing a weak positive relationship. D) Relationship between VMI score and the CBCL school competence score, showing a weak positive relationship. E) Relationship between the visual perception subtest score and the CBCL school competence score, showing a weak positive relationship. F) Relationship between the motor coordination subtest score and the CBCL school competence score, showing a weak positive relationship.

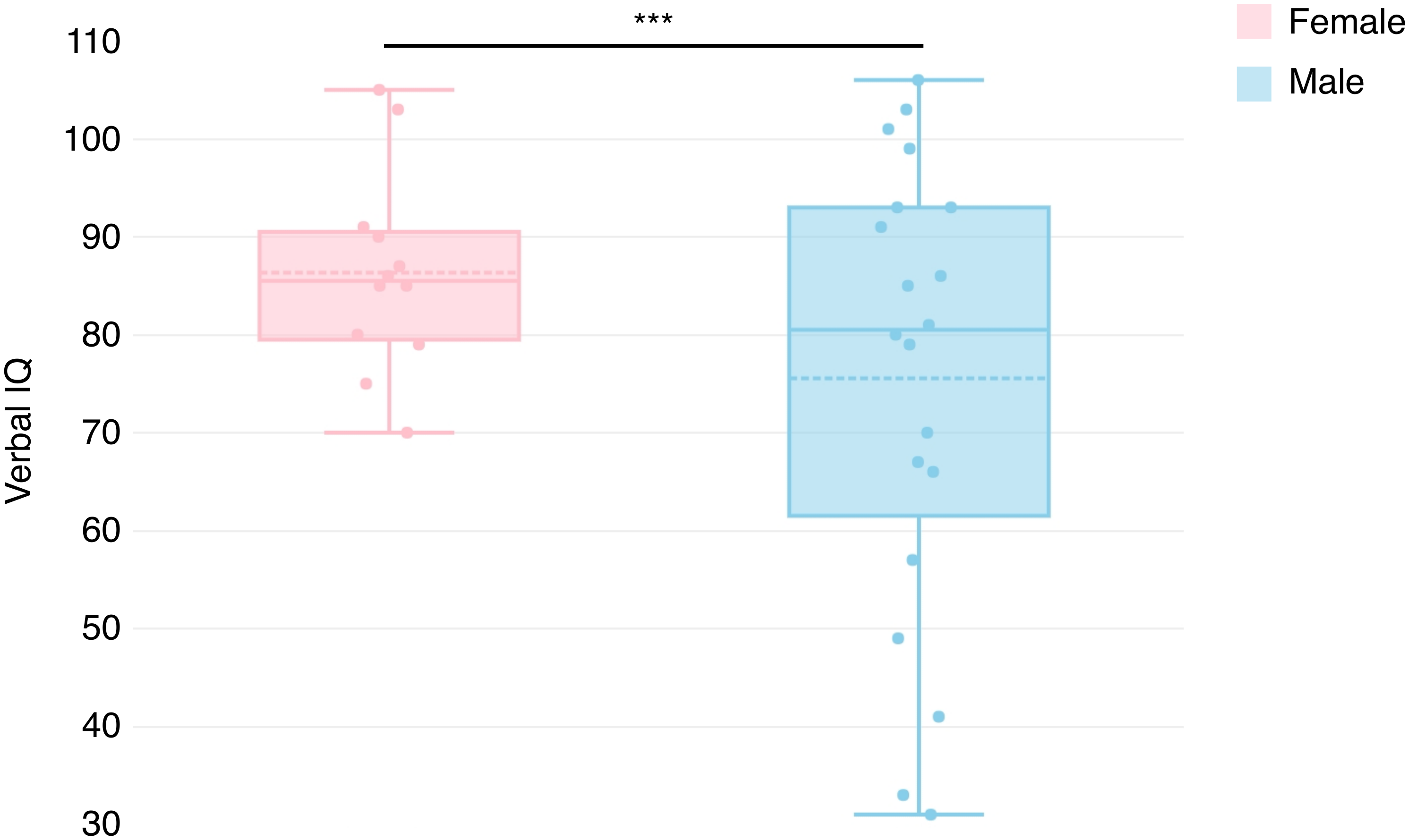

**Figure S7. Sex differences in verbal IQ.** Verbal IQ, showing that females (n = 12) score significantly higher than males (n = 20).

***, p < 0.001
